# Clonal phylogenies inferred from bulk, single cell, and spatial transcriptomic analysis of cancer

**DOI:** 10.1101/2023.02.26.530145

**Authors:** Andrew Erickson, Sandy Figiel, Timothy Rajakumar, Srinivasa Rao, Wencheng Yin, Dimitrios Doultsinos, Anette Magnussen, Reema Singh, Ninu Poulose, Richard J Bryant, Olivier Cussenot, Freddie C Hamdy, Dan Woodcock, Ian G Mills, Alastair D Lamb

**Author notes:** Corresponding Author: Dr Alastair D Lamb, CRUK Clinician Scientist Fellow & Honorary Consultant Urological Surgeon, Nuffield Department of Surgical Sciences, University of Oxford, Old Road Campus Research Building, Oxford OX3 7DQ.

## Abstract

Epithelial cancers are typically heterogeneous with primary prostate cancer being a typical example of histological and genomic variation. Prostate cancer is the second most common male cancer in western industrialized countries. Prior studies of primary prostate cancer tumor genetics revealed extensive inter and intra-patient tumor heterogeneity. Recent advances have enabled extensive single-cell and spatial transcriptomic profiling of tissue specimens. The ability to resolve accurate prostate cancer tumor phylogenies at high spatial resolution would provide tools to address questions in tumorigenesis, disease progression, and metastasis. Recent advances in machine learning have enabled the inference of ground-truth genomic single-nucleotide and copy number variant status from transcript data. The inferred SNV and CNV states can be used to resolve clonal phylogenies, however, it is still unknown how faithfully transcript-based tumor phylogenies reconstruct ground truth DNA-based tumor phylogenies. We sought to study the accuracy of inferred-transcript to recapitulate DNA-based tumor phylogenies.

We first performed in-silico comparisons of inferred and directly resolved SNV and CNV status, from single cancer cells, from three different cell lines. We found that inferred SNV phylogenies accurately recapitulate DNA phylogenies (entanglement = 0.097). We observed similar results in iCNV and CNV based phylogenies (entanglement = 0.11). Analysis of published prostate cancer DNA phylogenies and inferred CNV, SNV and transcript based phylogenies demonstrated phylogenetic concordance. Finally, a comparison of pseudo-bulked spatial transcriptomic data to adjacent sections with WGS data also demonstrated recapitulation of ground truth (entanglement = 0.35). These results suggest that transcript-based inferred phylogenies recapitulate conventional genomic phylogenies. Further work will need to be done to increase accuracy, genomic, and spatial resolution.

## Introduction

It is generally accepted that cancers develop and evolve by adaptive genetic and molecular changes over time (Nowell 1976; Greaves and Maley 2012; Black and McGranahan 2021). Sequential selection from this process of evolution leads to clones and subclones with altered phenotype leading to more aggressive behaviour. Ultimately, these phenotypic changes lead to metastatic spread and drug resistance, which is responsible for the majority of cancer-related deaths (Gupta and Massagué 2006).

It is necessary to distinguish accurately tumour heterogeneity and determine clonal evolution by identifying the clonal source of metastatic disease. This not only has an impact on the understanding of tumour progression but the relationship between clonal composition and the index lesion is also important and clinically relevant for both molecular diagnostics and focal therapy (Lamb et al. 2017; Reiter et al. 2019; Erickson et al. 2021). Indeed, it would help and support treatment decision-making by using new markers allowing to determine whether cells are indicative of aggressive disease or to predict sensitivity to treatment.

One of the challenges to understand the tumour heterogeneity is that origin of mutations occurring in cancer can be hereditary or somatic. Although identification of inherited mutations is relatively straightforward, these are only responsible for 5 to 10% of all cancer (Nagy et al. 2004; Garber and Offit 2005; Leon et al. 2021).By contrast, post-developmental somatic genetic alterations are usually only present in a small fraction of clonally-expanding cells but constitute the most common cause of cancer (Milholland et al. 2017). To identify these somatic mutations *in situ*, techniques such as laser capture microdissection have been employed, but this requires pre-knowledge to isolate a specific cell type or region of interest from a tissue section (Asp et al. 2020) and so limits the ability to undertake a *de novo* spatial clonal analysis. Recently, these limitations have been overcome by spatial transcriptomics, which allows the analysis of gene expression profiles in a tissue sample while preserving spatial tissue architecture. This approach captures transcripts *in situ*, with sequencing of barcoded reads carried out *ex situ* and then mapped back to the cells of origin (Larsson et al.2021; Ståhl et al. 2016). This cutting-edge technology permits visualisation and in-depth analysis of intra-tumoral heterogeneity and could permit spatial analysis of clonal evolution.

Clonal evolution and, more precisely, the relationship between clones and subclones is often represented and visualised by phylogenetic trees (Beerenwinkel et al. 2015; Schwartz and Schäffer 2017). These phylogenetic trees have been used mainly in recent years to study data derived from DNA sequencing (Schwartz and Schäffer 2017). However, to use spatial transcriptomics to study clonal evolution, it is necessary to know whether RNA can also be used to determine clonal phylogenetic hierarchies. In this meta-analysis, we investigate the correlation between DNA sequencing data and RNA sequencing data using phylogenies derived from inferred single-nucleotide variants (SNV) and copy-number variants (CNV) in order to determine whether transcriptome-derived phylogenies can accurately reflect genome-based phylogenies.

## Results

### Transcriptome and Genome Derived clonal phylogenies from single cancer cells

In order to benchmark performance of transcriptome-derived phylogenies, we first identified an individual cancer cell dataset with simultaneously isolated DNA and RNA (SIDR) from single cells (Han et al. 2018). The authors performed SIDR resulting in paired DNA and RNA nucleic acid extractions from isolated single cells of three different cancer cell lines: HCC827, MCF7 and SKBR3. They then performed whole-genome sequencing (WGS) and RNA-sequencing on the extracted nucleic acids. Given the cell purity, we hypothesized that WGS and RNA sequencing data from these individual cancer cells could be analyzed in an “in-silico” experiment to benchmark performance of transcriptome and genome-derived phylogenies.

We performed secondary analyses of the published, publicly available DNA and RNA sequencing data from Han et al (Han et al. 2018). After quality control (Han et al. 2018), we identified a total of 30 cells that had both sufficient quality DNA and RNA sequencing data, resulting in a dataset of a total of 10 MCF7 cells, 7 HCC827 cells, and 13 SKBR3 cells for analysis. We performed genomic SNV (gSNV) and inferred RNA-based SNV (iSNV) analyses from all cells, derived dendrograms, and performed tanglegram analysis to compare gSNV and iSNV dendrograms. In analysis of gSNVs and iSNVs, we observed a high concordance of transcriptome and genomic phylogenies (**Figure 1**, entanglement = 0.097). Next, we performed genomic CNV (gCNV) and inferred RNA-based CNV (iCNV) analyses from all cells, derived dendrograms, and performed tanglegram analysis to compare gCNV and iCNV dendrograms. In analysis of gCNVs and iCNVs, we also observed a high concordance of transcriptome and genomic phylogenies (**Figure 2**, entanglement = 0.11). We therefore concluded that RNA-derived inference of genomic SNVs and CNVs in three purified single cell populations generated strong phylogenetic concordance.

**Figure 1.**
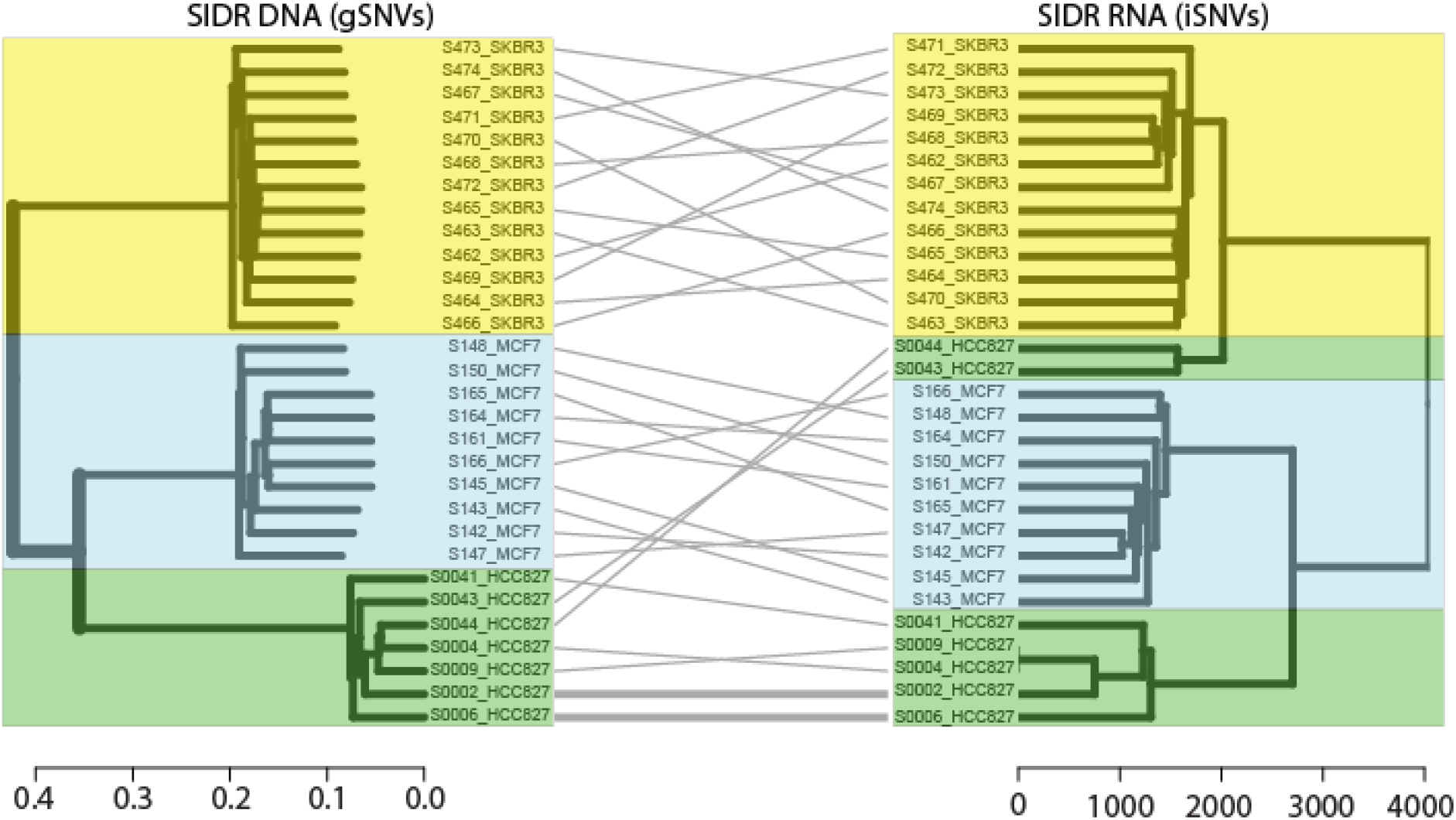
Comparison of in-silico clonal phylogenies from single tumour cells with co-isolated DNA and RNA (Han et al., Genome Res 2018). Dendrograms constructed from clustering of transcript-based inferred single-nucleotide variants (DENDRO) and ground truth DNA-based single-nucleotide variant calls (GATK) and compared by tanglegram. Colours correspond to individual cell lines (yellow: SKBR3, green: HCC827, and light blue: MCF7). Entanglement of the phylograms was 0.097 (an entanglement value of 1 corresponds with full entanglement of two phylograms, whereas an entanglement value of 0 corresponds with no entanglement).

**Figure 2.**
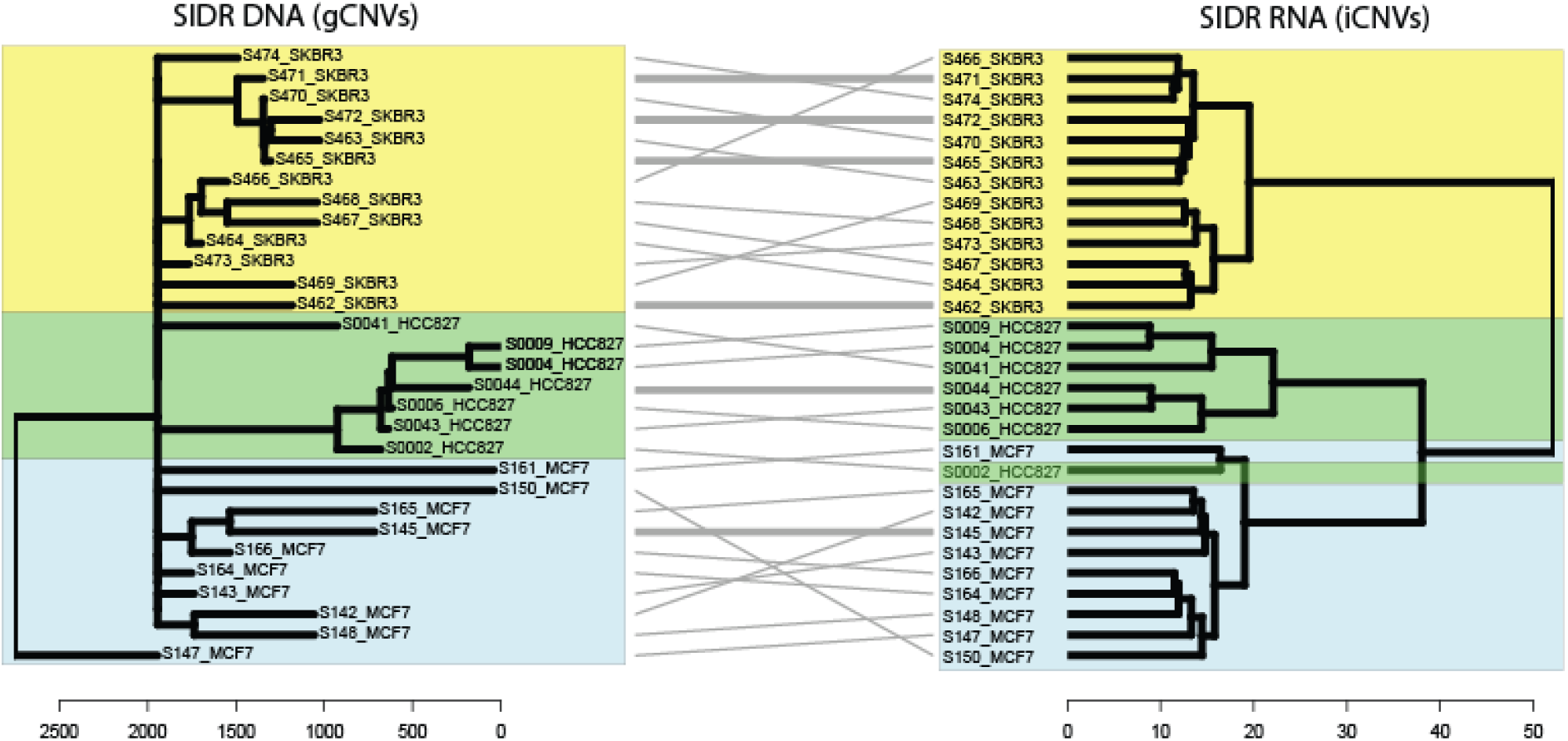
Comparison of in-silico clonal phylogenies from single tumour cells with co-isolated DNA and RNA (Han et al., Genome Res 2018). Dendrograms constructed from clustering of transcript-based inferred copy-number variants (inferCNV) and ground truth DNA-based copy number variant calls (WGS-Ginkgo) and compared by tanglegram. Colours correspond to individual cell lines (yellow: SKBR3, green: HCC827, and light blue: MCF7). Entanglement of the phylograms was 0.11 (an entanglement value of 1 corresponds with full entanglement of two phylograms, whereas an entanglement value of 0 corresponds with no entanglement). As adapted from Erickson et al., Nature, 2022, Extended Data Fig. 1a.

### Transcriptome and Genome Derived clonal phylogenies from bulk prostate cancer sequencing

Having established high *in-silico* concordance of transcriptome and genome-derived phylogenies, we then sought to study prostate cancer sequencing data from patients with paired DNA and RNA extracted from the same tumors. Gundem and colleagues reported WGS data from 55 disseminated tumor samples, from 10 patients that underwent rapid-autopsy after death due to prostate cancer (Gundem et al. 2015). A subset of n = 7 tumor specimens from patient A21 also underwent RNA-sequencing (Bova et al. 2016).

We performed secondary analyses of RNA sequencing data from Bova et al. and obtained iSNV and iCNV calls. From the iSNV and iCNV calls, we separately performed phylogenetic analyses through hierarchical clustering, resulting in iSNV and iCNV derived dendrograms (**Figure 3a**). In both iSNV and iCNV analyses, liver metastases (C, G, H, E) clustered together. In both iSNV and iCNV analyses, Clones F, A and J also clustered together. Clone I, clustered together with the liver metastases in iCNV analyses, but not in the iSNV analyses. Taken together, the iSNV and iCNV dendrograms reflect the manually assembled clonal phylogeny published by Gundem et al, (Gundem et al. 2015).

**Figure 3.**
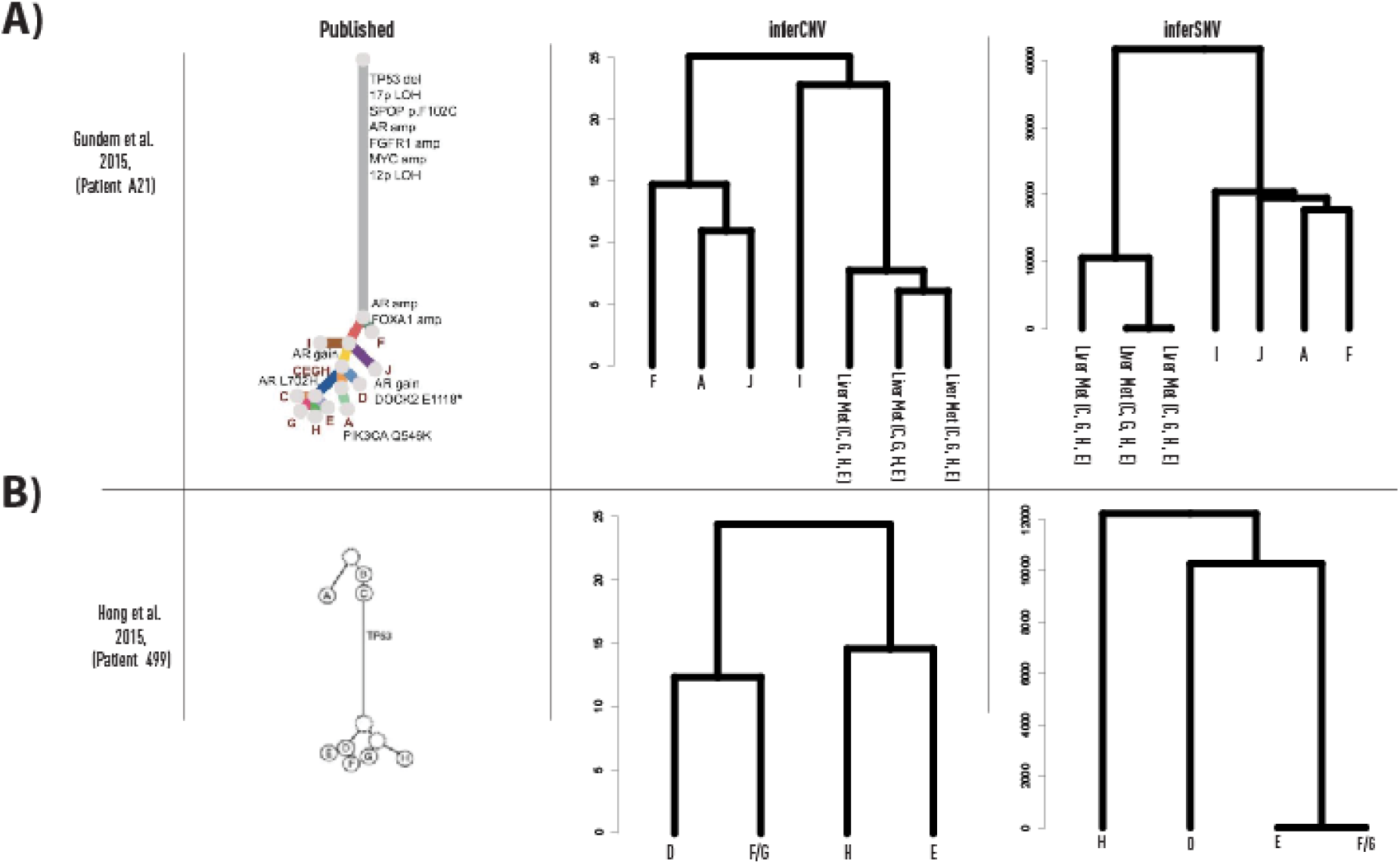
Comparison of published DNA-based prostate cancer clonal phylogenies and transcript-based inferred single-nucleotide and copy-number variant derived dendrograms. a) Phylogeny from patient A21, as published and reproduced from Gundem et al., Nature, 2015. Transcript data were available only for a subset of specimens. b, Phylogeny from patient 498, as published and reproduced from Hong et al., Nat. Comms, 2015. Transcript data available for a subset of specimens. inferCNV-based clonal phylogenies adapted from Erickson et al., Nature, 2022, Extended Data Fig. 1b.

Next, we analyzed data from patient 498, analyzed by Hong et al.(Hong et al. 2015). This patient’s primary prostate cancer progressed to distant skeletal metastases, which then further re-seeded the prostatic bed. Of the n = 7 reported specimens, a total of n = 4 also underwent RNA sequencing. We performed secondary analyses of the RNA sequencing data and obtained iSNV and iCNVcalls. From the iSNV and iCNV calls, we separately performed phylogenetic analyses through hierarchical clustering, resulting in iSNV and iCNV derived dendrograms (**Figure 3b**). In contrast to the results from Gundem et al., both iSNV and iCNV presenting differing tree patterns as compared to one another.

We then analyzed data from primary prostate cancer cases 6, 7 and 8, analyzed by Cooper et al., who each underwent radical prostatectomy, from which multiple tissue punches of both normal and tumor regions were sampled (Cooper et al. 2015). The samples then underwent WGS, which were subsequently analyzed and tumor phylogenies were manually produced. From a subset of the same specimens, adjacent tissue sections were taken and subjected to RNA microarray analysis. Additionally, each patient had a blood sample taken, that also underwent RNA microarray analysis. Being microarray data, we were unable to derive iSNV and iCNVs. Therefore, we built a custom pipeline to analyze and cluster the RNA microarray data directly, to generate hierarchical clustering represented as a dendrogram. To benchmark this pipeline, we first compared gCNV and gSNV to SIDR data (Supplementary Figure) and observed entanglement values of 0.21 and 0.16 respectively. Having established this pipeline, we then applied it to the microarray data from Cooper et al to generate dendrograms. These dendrograms were then analyzed in comparison to the published WGS-based gDNA phylogenies (**Figure 4**). In all three patients, the blood specimen clustered separately from the prostate tumor and normal tissue specimens. In cases 7 and 8, the (multiple) normal tissue specimens clustered together and distinctly clustered separately from the tumors, whereas in case 6 the two normals clustered with T2, T3 and T4, separate from T1. Taken together, RNA-microarray derived dendrograms were able to recapitulate manually assembled WGS-derived gDNA phylogenies.

**Figure 4.**
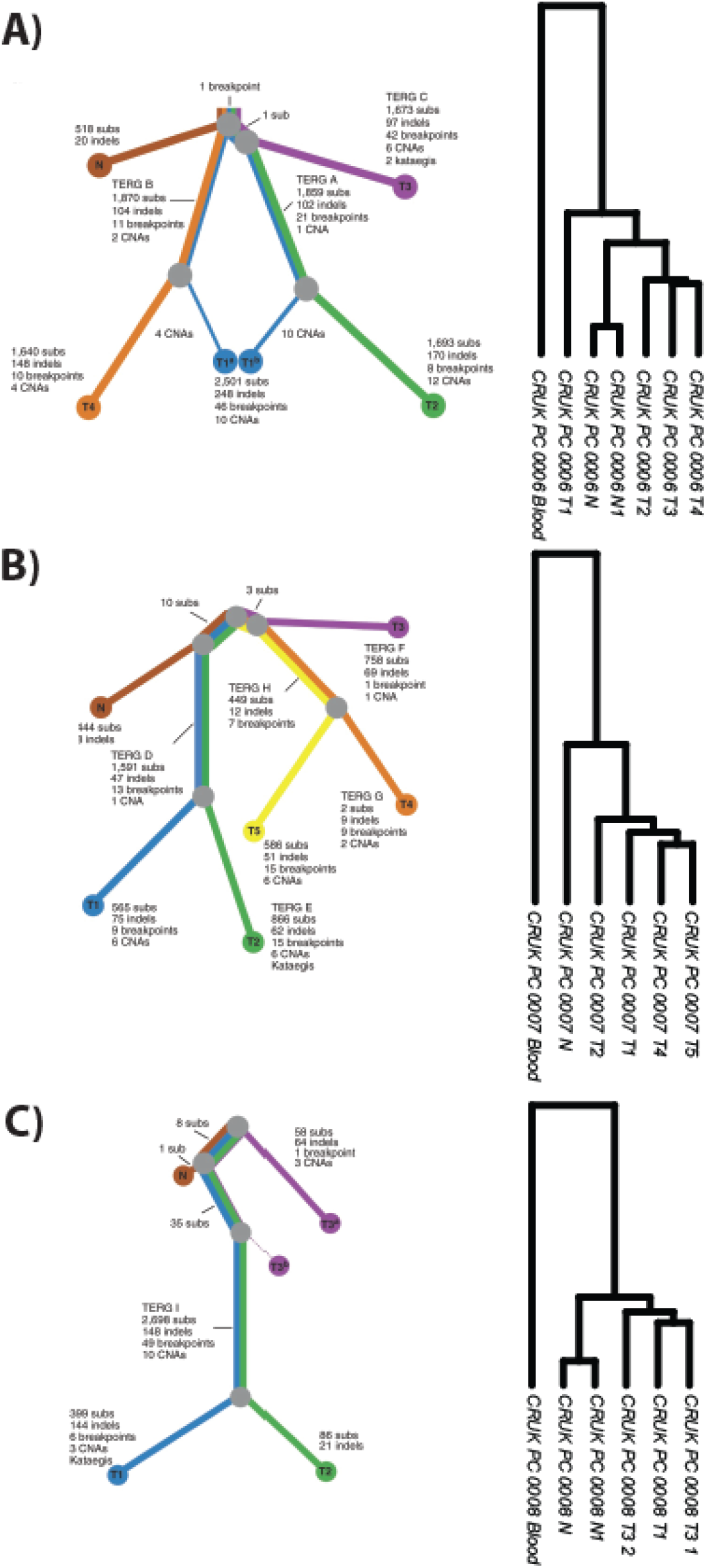
Comparison of published DNA-based (WGS) phylogenetic trees (left) as compared to novel RNA-based (RNA Microarray) phylogenies (right) from Cooper et al., 2015. A) Phylogenies from patient CRUK0006, B) Phylogenies from patient CRUK0007, C) Phylogenies from patient CRUK0008. RNA phylogenies include blood samples not presented in DNA-based phylogenetic trees.

**Supplementary Figure 1.**
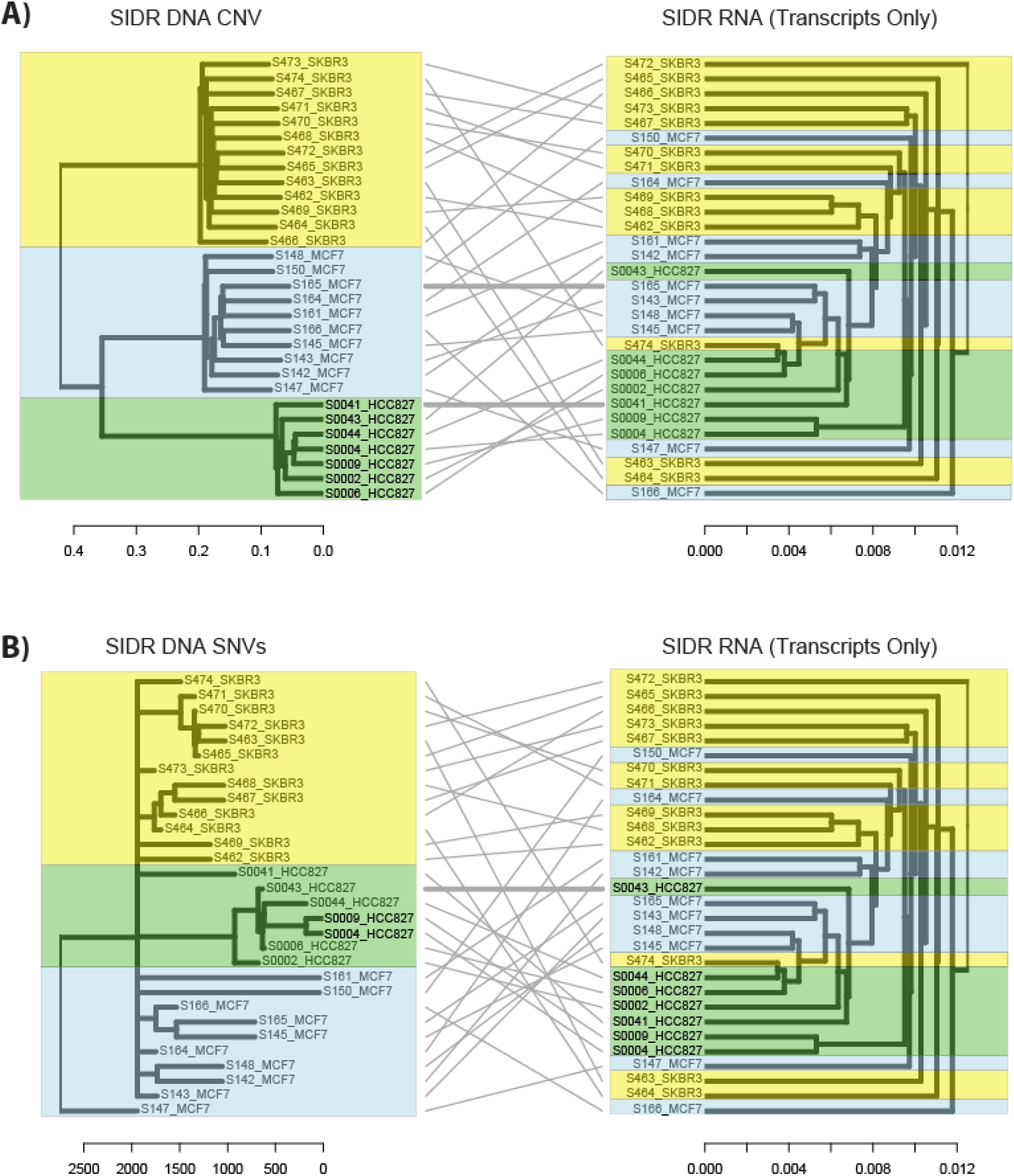
Comparison of in-silico clonal phylogenies from single tumour cells with co-isolated DNA and RNA (Han et al., Genome Res 2018). **A)** Dendrograms constructed from ground truth DNA-based copy number variant calls (WGS-Ginkgo) and direct transcripts (hierarchical clustering) and compared by tanglegram. Colours correspond to individual cell lines (yellow: SKBR3, green: HCC827, and light blue: MCF7). Entanglement of the phylograms was 0.21 (an entanglement value of 1 corresponds with full entanglement of two phylograms, whereas an entanglement value of 0 corresponds with no entanglement). **A)** Dendrograms constructed from ground truth DNA-based single-nucleotide variant calls (DENDRO) and direct transcripts (hierarchical clustering) and compared by tanglegram. Colours correspond to individual cell lines (yellow: SKBR3, green: HCC827, and light blue: MCF7). Entanglement of the phylograms was 0.16 (an entanglement value of 1 corresponds with full entanglement of two phylograms, whereas an entanglement value of 0 corresponds with no entanglement).

### Transcriptome and Genome Derived clonal phylogenies from bulk WGS and spatial transcriptomics from multi-region prostate cancer sequencing data

Next, we then sought to determine the ability of spatial transcriptome derived tumor phylogenies to recapitulate gDNA based phylogenies. Spatial transcriptomics generates transcriptome signale from poly-A captured short 3’ RNA sequences of up to 200 bp length, sufficient for hg38 alignment and, we deduced, sufficient to enable iCNV analysis. Berglund and colleagues performed spatial transcriptomics (ST) (Ståhl et al. 2016) on a total of n = 12 prostate tissue regions from a patient that underwent radical prostatectomy (Berglund et al.2018). Of these sections, a total of n = 4 were detected to have prostate cancer. The authors also performed WGS on adjacent serial sections from each of these 12 tissue sections, as well as a matched blood specimen from the same patient. Given that WGS is not spatially resolved, we performed ‘pseudo-bulked’ iCNV analyses on ST data from all 12 sections, and generated a clonal phylogeny in the form of a dendrogram. We also performed gDNA CNV calling from each of the 12 sections to generate a clonal phylogeny which was represented as a dendrogram. We then compared the iCNV and gCNV derived dendrograms using a tanglegram and observed a degree of concordance consistent with the resolution of the data (**Figure 5**, entanglement = 0.35). Interestingly, three of the tumor regions (P2_4, P1_3, P1_2) clustered together in the iCNV analysis, whereas they were represented on different subclusters in the gCNV phylogeny, suggesting that the iCNV approach may have generated a more accurate clustering in this case.

**Figure 5.**
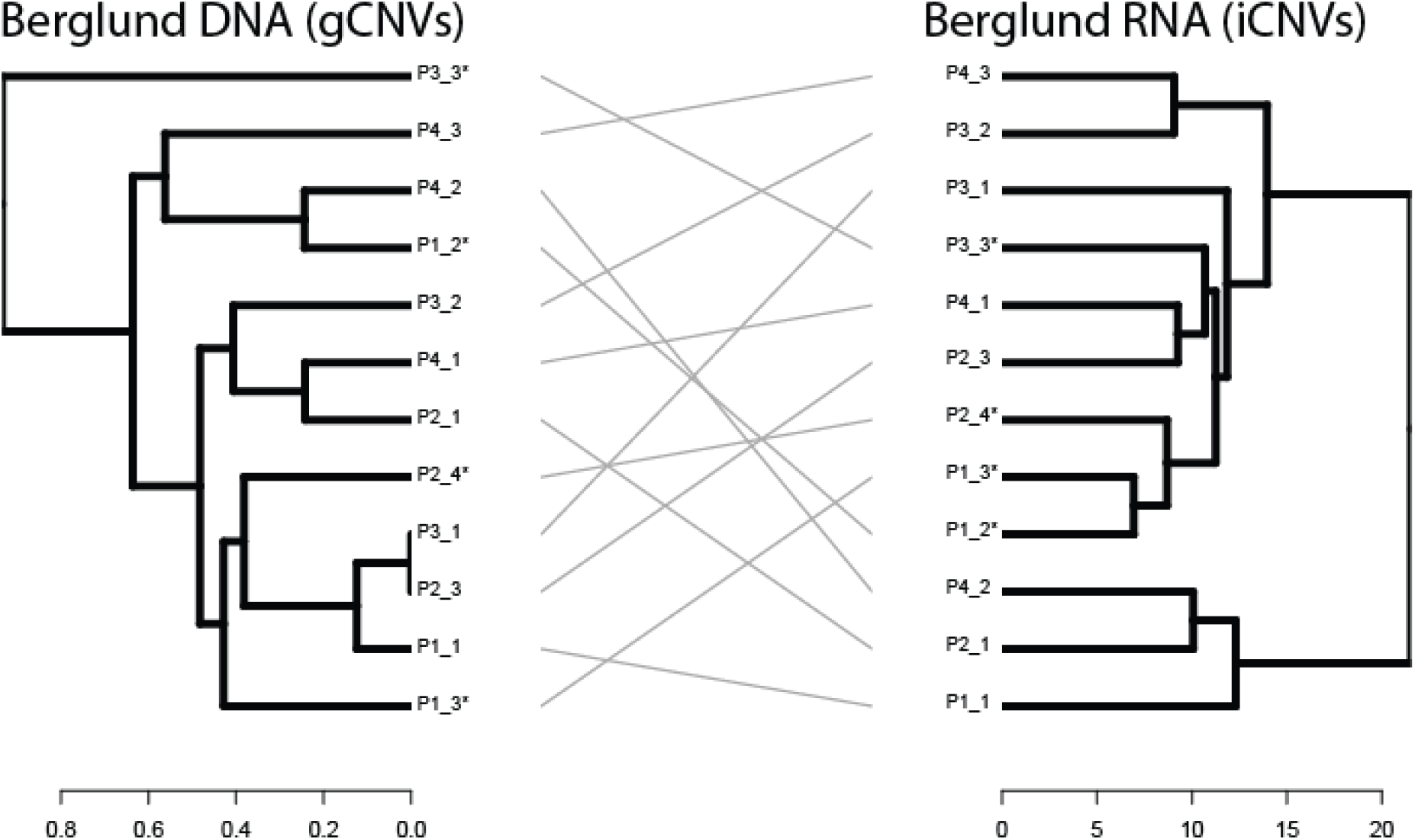
Comparison of DNA-based (WGS) phylogenetic trees (left) as compared to transcript-based inferCNV clonal phylogenies (right) from Berglund et al., 2018. DNA dendrogram constructed using patient-matched blood sample as a reference: such data were not available for inferCNV. Entanglement of the phylograms was 0.35 (an entanglement value of 1 corresponds with full entanglement of two phylograms, whereas an entanglement value of 0 corresponds with no entanglement). A label with the ending of * represents a section containing histologically detected cancer.

## Discussion

Results from single-cancer cells demonstrate that transcriptome-derived iCNV and iSNV phylogenies are highly concordant with ground truth gDNA based phylogenies. In our in-silico analyses, the analyzed data represent a highly selected and well controlled set of cells, with a 1:1 pairing of data resulting in extremely low entanglement values of the resultant tanglegrams. These results are in line with findings by Han et al., where they reported positive correlations for all three cell lines between gCNV and mRNA expression levels that were binned across the genome (Han et al. 2018). Our quantitative results in single-cells were supported by qualitative comparisons in prostate cancer cells where we did not have access to all ground truth data to enable a true like-to-like comparison.

There are limitations to consider in the construction of transcriptome-derived inferred phylogenies. First, the design and resolution of the genetic sequencing technologies can greatly affect the ‘resolved signal’. For example, only 2% of the entire genome is translated into proteins (International Human Genome Sequencing Consortium 2004), and thus the genomic coverage of the transcriptome represents a sub-fraction of potential data for mapping tumor phylogenies. This is further compounded by variable coverage within transcripts themselves: many modern scRNAseq and spatial transcriptomics techniques, such as Chromium and Visium offered by 10x Genomics, perform polyA capture, resulting in sequencing of 75-300 bp near the end of transcripts. Further, for iSNV approaches (Petti et al. 2019; Zhou et al. 2020), the coverage of transcribed SNV loci can be extremely low being confined to the exome. Potential issues with iSNVs seem to be mitigated in iCNV approaches (Patel et al. 2014; Gao et al. 2021; Elyanow et al. 2021), which incorporate machine learning algorithms to bin genomically adjacent transcripts. Additionally transcriptional regulation programs (Lee and Young 2013; Bradner et al. 2017; Davies et al. 2020) can affect transcription without any changes to copy-number status: these may result in false positive or negatives in iCNV analyses. Indeed, Han et al observed a discrepancy in Chromosome 3 gCNV calls and expression profiles (Han et al. 2018). Finally, one key factor affecting the ability of iCNV/iSNV (as well as gCNV and gSNV) approaches is use of well annotated references. All of the patient-derived WGS analyses in the data used in this publication had access to reference blood controls for calling gCNVs and gSNVs. Such data are not often taken or obtained for RNA sequencing, and thus are unavailable for iCNV and iSNV calling. This can also be further compounded by tissue or cell-of-origin transcriptional programs unrelated to copy-number alterations. Spatial transcriptomic data offers the opportunity to compensate for this through selection of histologically normal regions as control references.

As the tumor evolution community moves increasingly to single cell and spatial resolution, our ability to resolve clonal and subclonal tumor evolution patterns will greatly increase. Our results underscore the need for proper reference sets when calling iCNV and iSNV derived clonal phylogenies. These issues may be partly mitigated by next-generation iCNV and iSNV algorithms that incorporate both into combined iSNV+iCNV phylogenies (Gao et al. 2022). Other approaches incorporating evolutionary game theory through mathematical models could aid in resolving clonal phylogenies (Wölfl et al. 2022). Further work will also need to be done to identify and control for non copy-number alteration derived transcriptional regulation leading to further refinements in the ability of transcript-based clonal phylogenies to resolve ground truth.

## Methods

### Data Acquisition

In order to benchmark and validate methods to generate phylogenies derived from inferred single-nucleotide variants and copy-number variants, we reviewed the literature and found a recent publication which simultaneously extracted both DNA and RNA, from the same exact single tumor cells, and performed whole genome and whole transcriptome sequencing (Han et al. 2018). These public datasets contained data from 38 single cells that had been subject to simultaneous WGS and RNAseq using the SIDR methodology. Han et al describe a quality control process to determine which cells were satisfactorily sequenced for downstream analysis, leaving a total of 30 paired samples that passed all qc metrics.

Next, we reviewed the literature for publications and available data from patients with prostate cancer, who had both conventional bulk DNA and RNA sequencing applied to the same specimen, and from patients that had 3 or more total specimens. We identified patient A21 (Gundem et al. 2015; Bova et al. 2016), patient 498 (Hong et al. 2015). For further validation and comparison, WGS and RNA-microarray data were obtained from cases 6, 7 and 8 from Cooper et al. (Cooper et al. 2015).

Lastly, we obtained paired WGS sequencing data and paired Spatial Transcriptomics data from the n = 12 regions from a single patient in a recent publication (Berglund et al. 2018).

### Analysis of Single Cell Data

#### Quality Control of Single-Cell Whole Genome Sequencing Data

Only 38 paired cells were available with both scWGS and scRNAseq (Han et al. 2018). After removing the individual cells that failed either scWGS or scRNAseq QC left only 30 in common.

#### DNA Sequencing Preprocessing of Single-Cell Whole Genome Sequencing Data

Paired end sequencing data was aligned against the GRCh38 reference genome with the Burrow-Wheeler Aligner (0.7.17).

#### iSNV Calling from Single-Cell Whole Genome Sequencing Data

WGS variants were called using a pipeline broadly based on the GATK best practice Germline short variant discovery (SNPs + Indels) workflow using Picard (2.23.0) and GATK (4.1.7.0). This consisted of pre-processing the raw alignment to mark duplicate reads and perform base recalibration. Raw variants were called using GATK HaplotypeCaller in GVCF mode followed by GATK GenotypeGVCFs. Finally the raw variants were filtered to generate a downstream analysis ready cell by variant dataset.

The processed variants were converted to an Identity by State matrix, clustered and converted to dendrogram format in R using the SNPrelate package(Zheng et al. 2012)(Zheng et al.2017).

#### gCNV Calling from Single-Cell Whole Genome Sequencing Data

After preprocessing and QCing, n = 30 cells remained, and were then analyzed by Gingko (Garvin et al. 2015). BAM files were converted to .BED files using bamToBed in BedTools. We utilized a variable bin size of 50 kb, with 101 bp reads (Han et al. 2018). The clustering of CNV’s was performed using ward linkage and Euclidean distance as the distance metric. Copy-Number tree results were downloaded in Newick format for further downstream analysis.

#### RNA Sequencing Preprocessing of Single-Cell Whole Transcriptome Sequencing Data

Paired end sequencing data was aligned against the GRCh38 reference genome with STAR (2.7.3a) with per-sample 2-pass mapping and annotation with comprehensive gene annotation data from GENCODE GRCh38. Gene counts per cell were tabulated from aligned data using the featureCounts function from the Subread (1.6.4) package.

#### iSNV Calling from Single-Cell Whole Transcriptome Sequencing Data

iSNV calling from RNAseq data was performed according to the pipeline outlined by Zhou et al and based on GATK best practices (Zhou et al. 2020). The STAR aligned data underwent sorting, annotation with read group information, deduplication, SplitNCigarReads, realignment, and base recalibration, before variant calling with GATK (3.8.0) HaplotypeCaller. Raw iSNVs were processed by DENDRO to calculate a genetic divergence matrix between cells and to generate a phylogeny using hierarchical clustering (ward.D method).

#### iCNV Calling from Single-Cell Whole Transcriptome Sequencing Data

Data were analyzed using R version 4.0.1, and inferCNV (version 1.4.0) ([CSL STYLE ERROR: reference with no printed form.]). A merged file from the previously described pre-processing steps, containing feature counts for each cell, as well as a gene position file, and an annotation file were generated for input to inferCNV. An inferCNV object was created with no defined reference group. After creation of the InferCNV object, inferCNV was ran with the following parameters: cutoff = 0.1, cluster_by_groups = FALSE, denoise = TRUE, HMM = TRUE.

#### Comparison of Dendrograms from Single-Cells

For comparison of dendrograms created by WGS-CNVs (Gingko) and inferred CNV’s from RNA (InferCNV), the clust2.newick and infercnv.21_denoised.observations_dendrogram.txt files were imported into R and analyzed with packages *dendextend and phylogram.*

### Analysis of transcript derived phylogenies

RNA counts were analzyed, by comparing individual gene count values to the median (MED) and standard deviation (SD) values of global RNA count values per sample: if the count value was less than MED-SD, then it was assigned a value of −1, else if the count value was greater than MED+SD, then it was assigned a value of +1, else it was assigned 0. The resultant values from each sample or cell were converted into a phydat object using *phangorn*’s function phyDat(), with the parameters type=“USER”, levels = c(‘−1’, ‘0’, ‘1’). Pairwise distances between cells or tissue samples were calculated using the *phangorn* dist.ml() function with previously described phyDat() object as input. UPGMA clustering was applied using the *phangorn* upgma() function and converted to a dendrogram using the *dendextend* function as.dendrogram().

### Analysis of Spatial Transcriptomics Data

#### CNV Calling from Spatial Transcriptomics Data

Data were analyzed as previously described (Erickson et al. 2022) with the following exceptions. Original 1k array Spatial Transcriptomics data were obtained. As gCNV comparison data were from whole sections, all ST count data were ‘pseudo-bulked’ within sections, resulting in 12 pseudobulked count matrices for analyses. InferCNV was ran using standard parameters with no reference set. The resultant *infercnv.observations_dendrogram.txt* dendrogram was used for downstream tanglegram analysis.

#### Comparison of Dendrograms from WGS and ST

The original outputs for CNV calling from Berglund et al., were not available, and the ReadDepth package used to generate the calls has since been deprecated by the author (Miller). Thus, we ran a new pipeline using the WGS data from Berglund et al (Berglund et al.2018). FASTQ files were obtained and aligned to HG38. Battenberg CNV analyses (Nik-Zainal et al. 2012) were performed using the matched reference blood FASTQ data as the reference.

#### Copy number calling with Battenberg

The Battenberg package (v2.2.10) was used to determine copy number, and estimate tumour purity and ploidy from WGS data. Impute2 (v2.3.0) was used with GRCh38 loci for phasing germline heterozygous SNPs. The Battenberg pipeline was run with the following parameters: segmentation_gamma = 10, phasing_gamma = 10, platform_gamma = 1, min_ploidy = 1.6, max_ploidy = 4.8, min_rho = 0.13, max_rho = 1.02.

The *recal_subclones.txt* text files were downloaded for each of the 12 prostate tissues, and processed through a custom pipeline as follows. Battenberg CNV segments were binned into 1200 bp segments and aligned, generating n = 2439447 bins across the genome. CN amplifications and deletions were called at thresholded values of −1.5 and 2.5 respectively. Next, the processed bins from all samples were merged to create a CN bin matrix. CN calls for segments that were shared for all samples were dropped, resulting in a final matrix containing n = 28 discordant CN calls.

This CN matrix was then used similarly as described by Berglund et al., with the R package *pvclust*, and n = 1000 bootstraps. The structure of the cluster was converted to a dendrogram using the R package *dendrogram* for comparison to the inferCNV dendrogram via a tanglegram using the *dendextend* package (step2side).

## Data Access

Data from single cell experiments (Han et al. 2018) were previously deposited to ENA: PRJEB20144 (WGS) and PRJEB20143 (RNA). All sequence data from patient 499 (Hong et al. 2015) samples were previously deposited into the EGA Sequence Read Archive under accession number EGAS00001000942. RNA sequencing data from patient A21 (Bova et al.2016) were previously deposited into the EGA Sequence Read Archive under accession number EGAS00001001659. Sequencing data from patient 1 (Berglund et al. 2018) were previously deposited at the European Genome–Phenome Archive (EGA), hosted by the European Bioinformatics Institute (EBI), under the accession number EGAS0000100300.

## Competing Interest Statement

The authors have no conflicts of interest to declare.

## Acknowledgements

Computation used the Oxford Biomedical Research Computing (BMRC) facility, a joint development between the Wellcome Centre for Human Genetics and the Big Data Institute supported by Health Data Research UK and the NIHR Oxford Biomedical Research Centre. The views expressed are those of the author(s) and not necessarily those of the NHS, the NIHR or the Department of Health.

## Author contributions

A.E., A.L., and I.M. conceived the study. A.E., T.R., and S.R. performed computational experiments. All authors interpreted the data and wrote the manuscript.

## Notes

### Competing Interest Statement

The authors have declared no competing interest.

